# Selenium protects *Emiliania huxleyi* algae from a bacterial pathogen by modulating oxidative stress

**DOI:** 10.1101/2025.07.19.665712

**Authors:** Roni Beiralas, Shira Ben-Asher, Einat Segev

**Affiliations:** Department of Plant and Environmental Sciences, The Weizmann Institute of Science; Rehovot, Israel

## Abstract

Oxidative stress arises when cells fail to maintain redox balance, leading to the accumulation of reactive oxygen species (ROS) that can damage proteins, lipids, and DNA, causing cells to function poorly or die. In marine algae, oxidative stress is a hallmark of bacterial pathogenicity and usually appears before algae die. During the interaction of *Emiliania huxleyi* algae with *Phaeobacter inhibens* pathogenic bacteria, algae experience elevated levels of ROS that precede and likely drive cell death. Here, we tested whether antioxidants could improve algal survival by alleviating oxidative stress. Among several environmentally relevant antioxidants, we found that nanomolar concentrations of the trace metal selenium, in the form of H_2_SeO_3_, completely prevented algal death in co-cultures with *P. inhibens*. Measurements over time showed that selenium significantly lowered ROS levels in algal cells, reducing oxidative stress. This effect did not interfere with bacterial growth, suggesting that selenium acts by helping algae, not by harming bacteria. Our findings demonstrate that oxidative stress plays a central role in bacterial pathogenicity towards algae, and that selenium can protect algae by reducing this stress.

## Introduction

Algal blooms are vast oceanic events that are often terminated by a rapid demise (1). The bloom collapse is caused by various reasons, including viral infection and possibly pathogenic bacteria (2,3). In laboratory co-culture systems of algae and bacteria, bacterial pathogenicity towards algae has been well-documented over the past decade (3–8).

A unifying hallmark of bacterial pathogenesis is the intracellular accumulation of reactive oxygen species (ROS) in algal cells, a surge that typically precedes or coincides with algal death. Importantly, this oxidative stress does not necessarily originate from bacterial-produced ROS. Rather, bacterial pathogens often produce compounds that provoke the algal cells to generate ROS internally (3,4,7,9–11). In the model co-culture system of *Emiliania huxleyi* algae and *Phaeobacter inhibens* bacteria, abundant evidence support that bacterial metabolites or by-products induce oxidative stress within the algal cells, which subsequently triggers the demise of the algal population (3,9,10,12). In these co-cultures, most algal cells accumulate ROS, and ROS quenching rescues the algal population from collapsing (9). Transcriptomic data further show that genes involved in ROS detoxification are among the most strongly up-regulated algal transcripts during the *E. huxleyi* – *P. inhibens* interaction (3,10). Finally, a specific pathway of oxidative stress induction in algae was recently reported in the *E. huxleyi* – *P. inhibens* model system, demonstrating that algal-derived betaine stimulates bacterial H_2_O_2_ production, which in turn promotes algal death (12). Oxidative stress is central in various algal-bacterial interactions; ROS accumulation drives apoptotic death of bloom-formers such as *Phaeocystis globosa* and *Alexandrium tamarense* when exposed to *Microbulbifer* or *Brevibacterium* bacterial pathogens (4,7). The bacterial pathogen *Shewanella sp. IRI-160* which causes the death of several dinoflagellate species, was shown to produce a compound that increases intra- and extracellular ROS in dying algae (11). It appears that oxidative stress, driven by algal responses to bacterial signals or toxins, plays a central role in how bacteria cause algal death. Beyond promoting cell death, oxidative stress in microalgae has ecological consequences. Excess ROS promote the production of transparent exopolymer particles (TEP) and sticky mucilage (13–15). TEP enhances cell-cell aggregation, accelerates vertical carbon export, and can accumulate as dense foams that clog coastal infrastructure with detrimental influences on ecological systems (16–18). Oxidative stress is also linked to the synthesis and release of potent algal toxins: in *Microcystis aeruginosa*, exogenous ROS addition promotes microcystin production (15), while in several dinoflagellate species, oxidative stress is linked to algal toxicity (19,20). Thus, oxidative stress not only causes algal demise but also amplifies the ecological and economic impacts of bloom events through mucilage formation and toxin release. These harmful influences of oxidative stress highlight the need to explore strategies that can alleviate algal ROS accumulation and its downstream effects. For example, exogenous addition of antioxidants is already used in redox-management in higher plants. The addition of ascorbate, glutathione, or the trace metal selenium to the plant foliar or soil reduces ROS build-up and enhances crop yields (21,22).

In this study, we used the *E. huxleyi-P. inhibens* model system to test whether antioxidants can prevent bacteria-induced algal death by reducing oxidative stress. We screened a range of environmentally relevant antioxidants and found that nanomolar concentrations of selenium fully prevented algal death in co-culture with the bacteria. Time-resolved experiments using a ROS reporter showed that selenium significantly lowered oxidative stress levels in the algal population. These results suggest that selenium protects algae by reducing their oxidative stress response during interaction with bacteria. Our findings improve our understanding of algal-bacterial dynamics and point to low-dose micronutrient supplementation as a promising strategy for aquaculture and coastal bloom management.

## Results

### Algal oxidative stress increases in co-cultures with bacteria

To investigate oxidative stress in algae, we monitored *E. huxleyi* (also known as *Gephyrocapsa huxleyi* (23)) cultures grown either alone in mono-culture under axenic conditions or in co-culture with *P. inhibens* bacteria (see materials and methods). Over a 14-day incubation period, we tracked algal and bacterial growth using flow cytometry and plating, respectively. To assess oxidative stress in algal cells, we measured intracellular ROS using the fluorescent probe 2′,7′-dichlorodihydrofluorescein diacetate (H_2_DCFDA; see materials and methods) (Fig. 1). The fluorescence signal was detected via flow cytometry, which enabled us to quantify ROS-positive events specifically within the algal population. Algal death occurred in the co-cultures between days 7 and 10, while the axenic algal cultures remained stationary and did not exhibit a similar demise (Fig. 1A). Axenic cultures exhibited increased levels of intracellular ROS during the stationary phase (particularly at days 10 and 14), while algae in co-cultures showed significantly higher ROS levels at the same time (Fig. 1B). Notably, the elevated algal oxidative stress in the co-cultures compared with axenic cultures was significant on days 7, 10 and 14 – both prior to and during the algal death. Interestingly, the most pronounced difference in algal oxidative stress between co-cultures and axenic cultures occurred on day 7, before the onset of algal death (Fig. 1A and B). In all co-cultures, bacteria grew and reached the stationary phase on day 10 (fig. S1, dark gray line).

**Figure 1.**
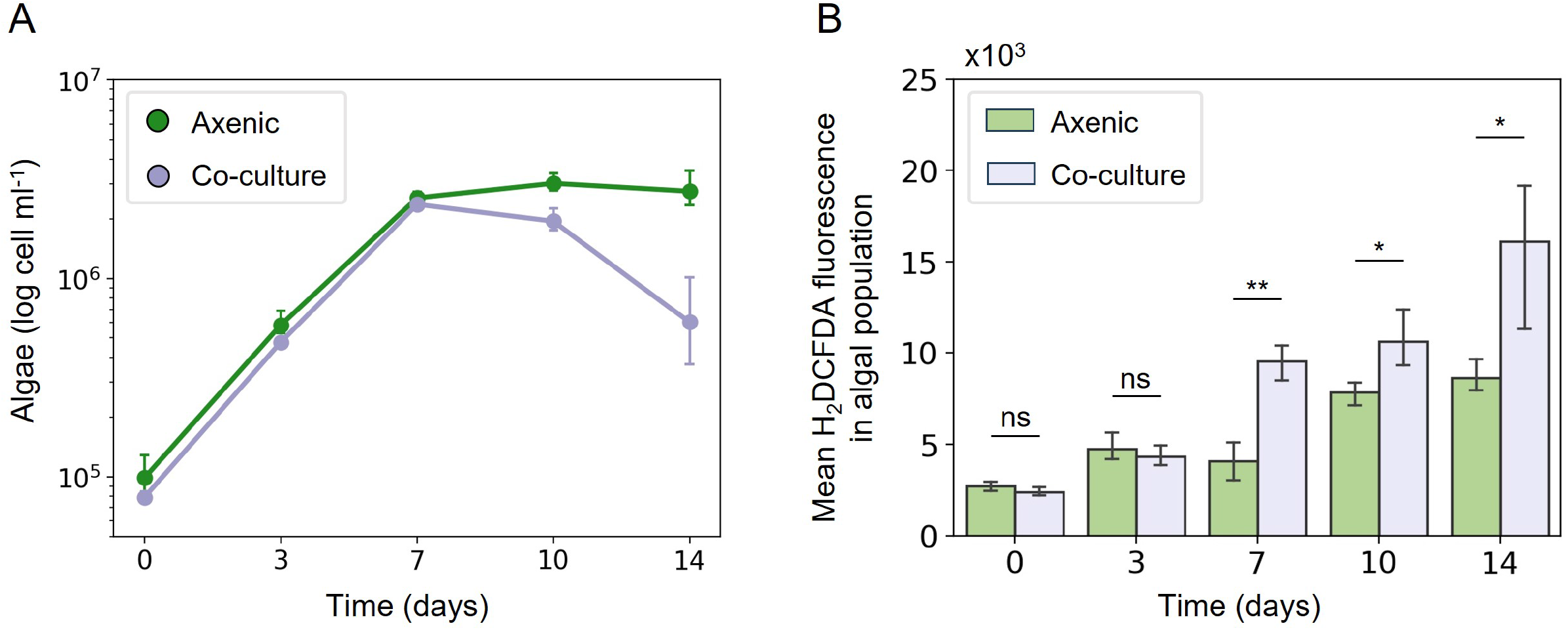
Algal oxidative stress in axenic cultures and co-culture with bacteria. (A) The growth of *E. huxleyi* CCMP2090 algae during 14 days, cultured axenically (green line) or in co-culture with *Phaeobacter inhibens* DSM17395 bacteria (purple line). (B) Mean fluorescence intensity of the probe 2’,7’-dichlorodihydrofluorescein diacetate (H_2_DCFDA) which indicates oxidative stress in *E. huxleyi* populations during 14 days of growth. Green bars – axenic algae, purple bars – algae grown with *P. inhibens* bacteria in co-culture. Statistical significance was calculated using a two-sample t-test to compare the fluorescent signal between axenic cultures and co-cultures. * *P* < 0.05, ** *P* < 0.005, ns – not significant. Each data point consists of 3 biological replicates, error bars designate ± SD.

Together, these findings reveal a correlation between oxidative stress and algal cell death. Notably, the rise in oxidative stress precedes the onset of algal death in co-cultures, suggesting that the oxidative stress induced by bacteria may contribute to the initiation of algal demise.

### Selenium prevents algal death in co-cultures with bacteria

Building on our observation that oxidative stress may play a role in bacterially mediated algal death, we investigated whether antioxidants could alleviate algal demise by reducing oxidative stress. To explore this, we screened various antioxidant compounds that are naturally found in the marine environment or produced by marine organisms, including selenium (24), ascorbate (25), dimethylsulfoniopropionate (DMSP) (26), and α-tocopherol (25). To evaluate the impact of these antioxidants, *E. huxleyi* algae were cultured both axenically and in co-culture with *P. inhibens* bacteria, and algal survival was monitored (Fig. 2).

**Figure 2.**
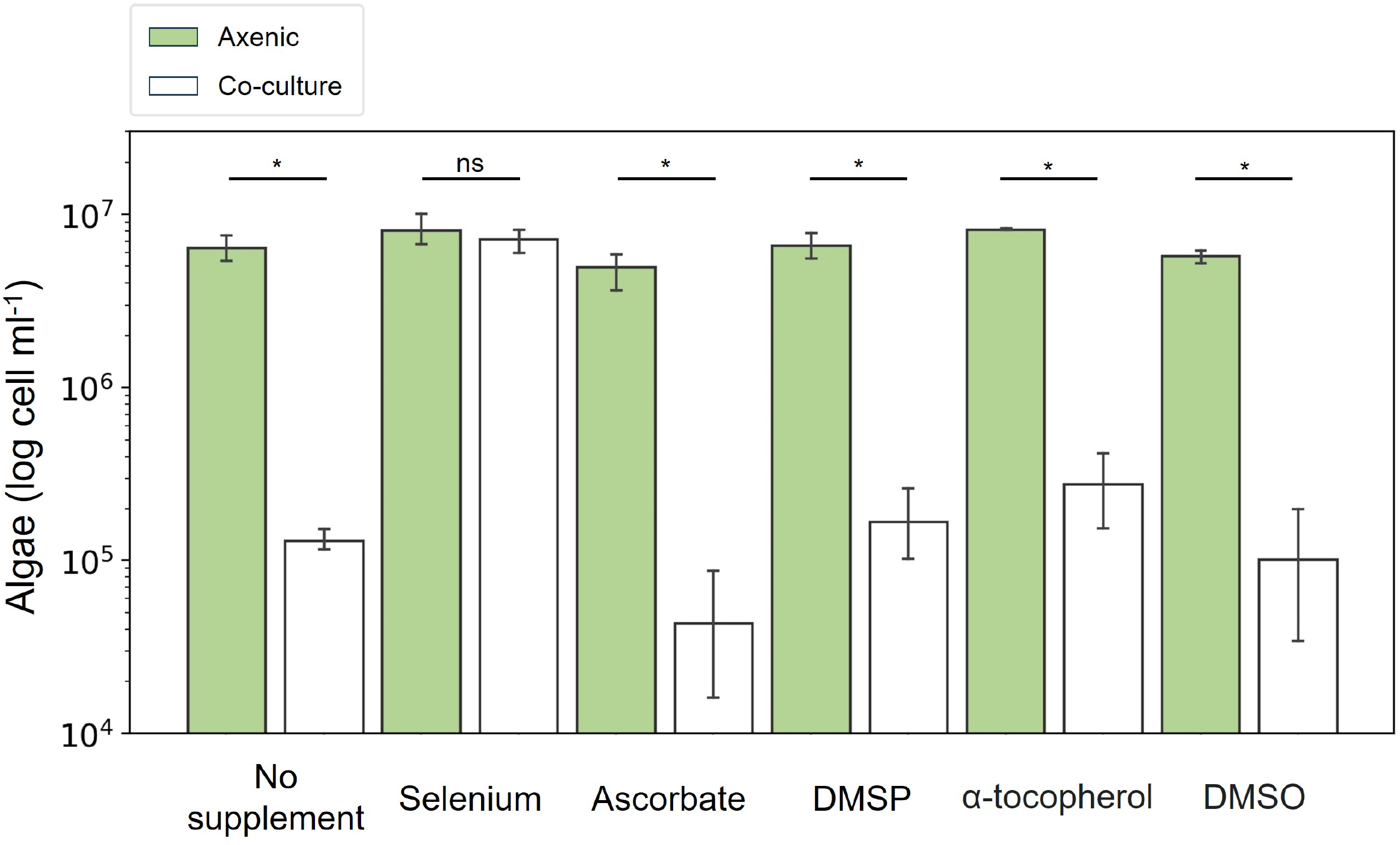
Algal viability in axenic and co-cultures supplemented with different antioxidant molecules. The algae *E. huxleyi* CCMP2090 were grown either axenically (green bars) or with *P. inhibens* DSM17395 bacteria (white bars). The cultures were supplemented on the day of algal inoculum (day -4) with different antioxidants – selenium (H_2_SeO_3_, 10 nM), ascorbate (L-ascorbic acid, 20 µM), DMSP (dimethylsulfoniopropionate, 20 µM) and α-tocopherol (20 µM). DMSO (dimethylsulfoxide, 60 µl) was used as the solvent of α-tocopherol hence it was supplemented to cultures as a control. Algal cell concentrations were measured at a single time point on day 17 of growth to account for the influence of the different antioxidants tested. Statistical significance was calculated using a two-sample t-test to compare the cell numbers between axenic cultures and co-cultures. * *P* < 0.005, ns – not significant. Each data point consists of 3 biological replicates, error bars designate ± SD.

Our results show that among the antioxidants tested, only selenium effectively prevented algal death in co-culture with *P. inhibens* bacteria (Fig. 2, fig. S2A, B). Algae in co-cultures treated with selenium remained at levels comparable to those in axenic cultures, indicating that selenium protected the algae from the pathogenic effects of bacteria. In contrast, ascorbate, DMSP, and α-tocopherol did not prevent algal death (Fig. 2, fig. S2A, B). Interestingly, the DMSO solvent of α-tocopherol and α-tocopherol itself delayed the onset and progression of algal growth and death in both axenic and co-culture conditions (fig. S2A, B). It is important to note that the initial algal inoculum on day -4 was identical across all treatments (see material and methods); the cell concentrations measured at day 0 represent algal densities after 4 days of growth under the various treatments. Despite the observed delay under DMSO and α-tocopherol treatment, algal collapse still occurred in co-cultures, although later, on day 17 instead of day 14.

Importantly, during the antioxidant screen, the growth of *P. inhibens* was monitored to ensure that the observed mitigation of pathogenicity was not due to inhibition of bacterial growth. Under all treatments, bacteria grew to comparable final densities (fig. S2C). Although ascorbate and α-tocopherol treatments resulted in slower bacterial growth rates, the bacterial populations ultimately reached the same final density as under control conditions, and induced algal death (Fig. 2, fig. S2B, C). Notably, *P. inhibens* bacteria in co-cultures supplemented with selenium grew at the same rate as in the control conditions, although the typical algal death was not observed (Fig. 2, fig. S2B, C).

These findings show that selenium prevents bacterial-induced algal mortality. Given its antioxidant properties, we next investigated whether selenium counters the pathogenic effects of *P. inhibens* in co-culture by lowering algal oxidative stress.

### Selenium reduces algal oxidative stress in co-cultures

Given the observed correlation between oxidative stress and algal death in co-cultures with bacteria, and the protective effect of selenium, we tested whether selenium prevents algal death by lowering oxidative stress in algal cells. We therefore treated with selenium both axenic *E. huxleyi* cultures and co-cultures with *P. inhibens* bacteria, then measured algal intracellular ROS levels using the H_2_DCFDA fluorescence probe. In parallel, we monitored algal and bacterial cell numbers to relate oxidative stress dynamics to growth patterns.

In the axenic cultures, algae remained viable regardless of selenium supplementation (Fig. 3A). Interestingly, axenic algae achieved significantly higher cell densities when supplemented with selenium compared to cultures without selenium starting from day 3 (Fig. 3A). In axenic cultures, oxidative stress levels were significantly lower in selenium-treated samples at all time points after day 0. While cultures without selenium showed a gradual increase in oxidative stress over time, selenium-treated cultures maintained consistently low levels throughout the experiment (Fig. 3B).

**Figure 3.**
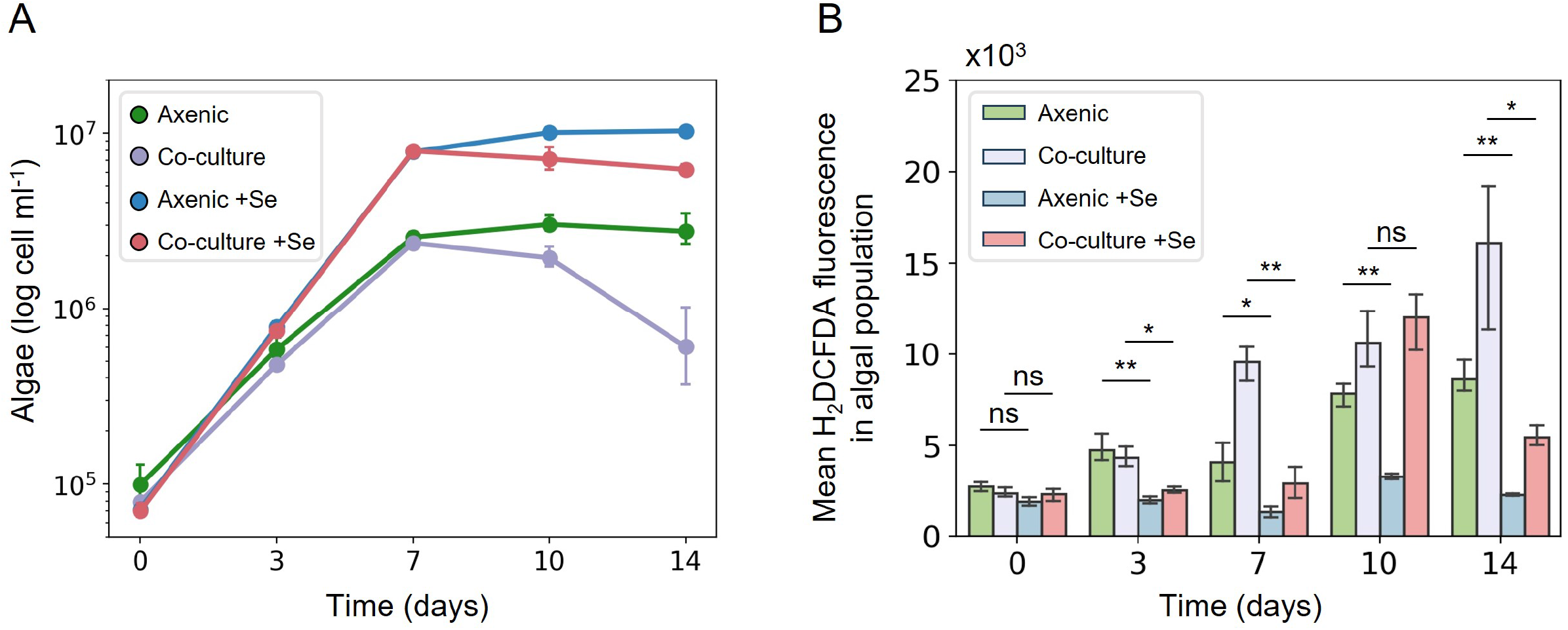
Oxidative stress dynamics in algal populations grown with and without selenium. (A) The growth of *E. huxleyi* CCMP2090 algae during 14 days cultured axenically, untreated and treated (+Se) with selenium (green and blue lines, respectively) or in co-cultures with *P. inhibens* DSM17395 bacteria, untreated and treated (+Se) with selenium (purple and pink lines, respectively). Statistical significance was calculated using a two-sample t-test to compare algal cell numbers between axenic cultures untreated and treated with selenium, and co-cultures untreated and treated with selenium. Starting from day 3, all comparisons resulted in significantly higher cell numbers in selenium-treated cultures with *P* < 0.05 on day 3 and *P* < 0.0005 on days 7, 10 and 14. (B) Mean fluorescence intensity of the probe 2’,7’-dichlorodihydrofluorescein diacetate (H_2_DCFDA) which indicates oxidative stress in *E. huxleyi* populations during 14 days of growth. Green and blue bars – axenic algae untreated and treated (+Se) with selenium, respectively. Purple and pink bars – algae grown with the bacterium *P. inhibens* untreated and treated (+Se) with selenium, respectively. Statistical significance was calculated using a two-sample t-test to compare the fluorescence signal between axenic cultures untreated and treated with selenium, and co-cultures untreated and treated with selenium. * *P* < 0.05, ** *P* < 0.005, ns – not significant. Each data point consists of 3 biological replicates, error bars designate ± SD.

In co-cultures with *P. inhibens* bacteria, selenium treatment preserved algal viability, in contrast to the rapid algal death observed in untreated co-cultures (Fig. 2, Fig. 3A). As in axenic cultures, selenium-treated co-cultures maintained significantly higher algal cell densities compared to untreated controls starting from day 3 (Fig. 3A). ROS measurements revealed significantly higher algal oxidative stress in untreated co-cultures on all days except day 0 and day 10 (Fig. 3B). While algal oxidative stress levels in untreated co-cultures increased steadily over time, co-cultures treated with selenium exhibited largely stable algal ROS levels, with a transient peak on day 10, followed by a marked decrease by day 14. These observations suggest that although algal oxidative stress still occurs in selenium-treated co-cultures, its reduced levels are not sufficient to promote algal death. Importantly, bacterial growth dynamics remained comparable between untreated and selenium-treated co-cultures, indicating that the protective effect of selenium on algae is not due to changes in bacterial growth (fig. S1, light gray line). These findings strongly support the conclusion that the protective influence of selenium stems specifically from its ability to reduce bacterial-induced oxidative stress in algae.

Taken together, our results show that oxidative stress is a central mechanism through which *P. inhibens* exerts pathogenic effects on *E. huxleyi*, and that selenium effectively reduces oxidative stress in the algal population. By preventing oxidative stress and mitigating bacterial pathogenicity, selenium presents a promising avenue for future applications.

## Discussion

### Oxidative stress is a hallmark of bacterial pathogenicity towards algae

Our data show that oxidative stress is an early and reproducible signature of the pathogenic interaction of *P. inhibens* bacteria and *E. huxleyi* algae. Algal intracellular ROS accumulate to significantly higher levels in co-cultures relative to axenic controls by day 7, well before any measurable loss of algal viability is detected (Fig. 1). This timing highlights the intracellular ROS level as a potential driver of algal mortality rather than a by-product of dying cells, similar to earlier reports in other bloom-formers exposed to algicidal bacteria (4,7,11).

### Selenium mitigates bacterial pathogenicity by alleviating algal ROS

Supplementing nanomolar concentrations of selenium prevented the algal population collapse even though *P. inhibens* bacteria reached identical densities in treated and untreated cultures (Fig. 2, Fig. 3, fig. S1, fig. S2). These observations suggest that algal protection was not the result of impaired bacterial growth. Selenium-treated algae in co-cultures accumulated significantly less ROS compared to untreated algae in co-cultures and remained viable even when bacteria reached their maximal density (Fig. 3, fig. S1). These results suggest that *P. inhibens* promotes algal death by altering the algal redox balance, possibly beyond a critical threshold. Chemically potent antioxidants such as ascorbate, DMSP and α-tocopherol failed to protect the algae, whereas selenium which is an essential trace element that cells incorporate as the amino acid selenocysteine (27), prevented algal death. The molecular mechanism that underlies the protective impact of selenium is currently unknow, but it likely acts by strengthening the catalytic antioxidant defenses of algae. In coccolithophores, selenocysteine is inserted into a large suite of selenoproteins, most notably glutathione peroxidases and thioredoxin reductases (27). *E. huxleyi* algae possess an unusually large selenoproteome, including multiple glutathione peroxidases and thioredoxin reductases (27–29). These enzymes, which depend on nanomolar concentrations of selenium for maximal activity, act as antioxidants with orders-of-magnitude higher efficiency than their cysteine-containing homologues (27). The addition of selenium could therefore boost the activity of this enzymatic arsenal, maintain ROS levels below lethal concentrations, and prevent the cell-death cascade that is triggered by pathogenic bacteria.

### Algal redox as a mortality predictor

Reducing oxidative stress, rather than suppressing the pathogen itself, appears to be sufficient to prevent algal mortality. Therefore, algal intracellular ROS could serve as a predictor of algal fate during this algal-bacterial interaction. Comparable redox signatures were reported in various algal-bacterial systems, where diverse bacterial metabolites promoted oxidative bursts that resulted in algal death (3,4,7,9–11). Viruses also exploit a similar algal redox vulnerability; infection of *E. huxleyi* with its EhV virus leads to elevated algal ROS early in the replication cycle of the virus, and chemical ROS quenching impairs virion production (30). Furthermore, EhV infection triggers a transient antioxidant response in the algal host, but this defense is insufficient, and the infected cells ultimately lyse (31). These observations highlight the centrality of host redox balance in interactions with diverse microbial antagonists.

### Host-microbe redox dynamics

The host redox balance shapes a variety of host-microbe interactions across kingdoms: in both mammalian and plant hosts, microbial metabolites and secreted effectors can promote substantial ROS accumulation in the host, and this oxidative surge is a major driver of tissue damage and inflammatory pathology (32–34). Whether such damage progresses or is contained depends on how effectively the host or the surrounding environment increases their antioxidant capacity. When detoxification networks are reinforced, either endogenously or through supplementation of micronutrients or antioxidants, the redox imbalance is reduced and disease severity declines (35,36). Our findings therefore place the *E. huxleyi-P. inhibens* interaction within a spectrum of redox-driven antagonisms and demonstrate that trace-elements can influence the outcome of these interactions.

### Environmental outlook

In the marine environment, dissolved selenium is patchy, enriched in dust-fed surface layers yet scarce in oligotrophic oceans (37). Regional pulses of selenium might influence the ability of *E. huxleyi* cells and blooms to resist or succumb to bacterial pathogens, or oxidative stress in general. To fully understand the cellular mechanisms of algal protection mediated by selenium, it will be important to integrate molecular insights with field-based studies, such as mesocosm experiments (38), in which selenium levels are manipulated within natural microbial communities. This comprehensive approach could open exciting avenues for evaluating micronutrient supplementation as a practical strategy in aquaculture and bloom management.

## Materials and methods

### Strains and growth conditions

The algal strain of *Emiliania huxleyi* CCMP2090 was purchased as an axenic culture from the National Center for Marine Algae and Microbiota (Bigelow Laboratory for Ocean Sciences, ME, USA). Algae were maintained in artificial seawater medium (ASW) prepared according to Sperfeld, *et al*. (39), and supplemented with L1 nutrients (NaNO_3_, 882 μM; NaH_2_PO_4_·2H_2_O, 36.22 μM), f/2 vitamins (thiamine HCl, 100 μg l^™1^; biotin, 0.5 μg l^−1^; vitamin B_12_, 0.5 μg l^−1^) and f/2 trace metals (Na_2_EDTA·2H_2_O, 4.36 mg l^−1^; FeCl_3_·6H_2_O, 3.15 mg l^−1^; MnCl_2_·4H_2_O, 178.1 μg l^−1^; ZnSO_4_·7H_2_O, 23 μg l^−1^; CoCl_2_·6H_2_O, 11.9 μg l^−1^; CuSO_4_·5H_2_O, 2.5 μg l^−1^; Na_2_MoO_4_·2H_2_O, 19.9 μg l^−1^). Stock solutions of nutrients, vitamins and trace metals were purchased from the Bigelow Laboratory. Algae were grown in standing cultures in a growth room at 18°C under a light/dark cycle of 16/8 hr. Illumination intensity during the light period was 130 μmol photons m−2 s−1. Absence of bacteria in axenic algal cultures was monitored weekly both by plating on ½ YTSS plates (containing yeast extract, 2 g l^−1^; tryptone, 1.25 g l^−1^; sea salts, 20 g l^−^1) and under the microscope.

The bacterial strain of *Phaeobacter inhibens* DSM 17395 was purchased from the German collection of microorganisms and cell cultures (DSMZ, Braunschweig, Germany). Bacteria were cultured in ½ YTSS medium, either in liquid culture or on agar plates, and incubated at 30 °C with shaking at 130 rpm.

### Algal cell counts

Algal growth was monitored using a CellStream CS-100496 flow cytometer (Merck, Darmstadt, Germany), with excitation at 561 nm and emission detection at 702 nm. For each sample, 50,000 events were recorded. Algal cells were gated according to event size and fluorescence intensity.

### Algal-bacterial co-culturing

Co-cultures of *E. huxleyi* and *P. inhibens* were established as follows: algal cells from a late-exponential phase culture were quantified using flow cytometry as described above. An inoculum of 10^3^ algal cells was introduced into 30 ml of ASW medium supplemented with nutrients, vitamins and trace metals. The time point of algal inoculation was designated ‘day -4’. After four days of algal growth, *P. inhibens* bacteria were prepared for co-culturing. Bacteria were harvested from 48 hr liquid culture in ½ YTSS medium, washed three times with ASW and diluted to an OD_600_ of 0.01. This suspension was further diluted to 1:1,000, and 20 µl was added to the algal cultures. The time point of bacterial addition into algal cultures was designated ‘day 0’. Co-cultures were incubated in the same growth conditions as described for axenic algal cultures. Sampling days are indicated relative to the time of bacterial addition.

### Monitoring bacterial growth in co-cultures

Bacterial abundance in co-cultures was assessed at various time points, as indicated. Samples were serially diluted in ASW and plated on ½ YTSS agar plates. Colony-forming units (CFUs) were counted after incubation, and bacterial concentrations in the original samples were calculated accordingly.

### Intracellular reactive oxygen species (ROS) measurements in algal cells

Intracellular ROS levels in algal cells were measured using the fluorescent dye 2′,7′-dichlorodihydrofluorescein diacetate (H_2_DCFDA; ThermoFisher), dissolved in DMSO. Fluorescence was detected using a CellStream CS-100496 flow cytometer, with excitation at 488 nm and emission collected at 528/46 nm. Algal cells were gated as described above for algal cell counts.

For each measurement, 400 µl of culture was sampled and split into two 200 µl sub-samples. One sub-sample served as a negative control and was treated with DMSO; the other was treated with 0.5 µM H_2_DCFDA. Samples were incubated in the dark for 30 minutes prior to measurement. Mean fluorescence intensity of the gated algal population was quantified using CellStreamAnalysis software. Background fluorescence (from the DMSO-treated control) was subtracted from the fluorescence of the H_2_DCFDA-treated sample to obtain the corrected intracellular ROS signal in the algal population.

### Antioxidants screen

Algal-bacterial co-cultures and axenic algal cultures were prepared as described above, with the addition of selected antioxidants at the time of algal inoculation (day -4). The antioxidants tested included 10 nM selenious acid (H_2_SeO_3_), 20 µM L-ascorbic acid (ascorbate), 20 µM dimethylsulfoniopropionate (DMSP), and 20 µM α-tocopherol. All compounds were dissolved in ASW, except α-tocopherol, which was dissolved in DMSO. To control for potential solvent effects, DMSO was added to control cultures. Algal and bacterial cell concentrations were monitored throughout the experiment as described above.

## Acknowledgements

We are grateful to Dr. Or Eliason for his valuable guidance in establishing the H_2_DCFDA fluorescence assay for quantifying intracellular ROS in the algal population. We thank all members of our lab for valuable discussions.

## Data availability statement

Data from this study is available from the corresponding author upon reasonable request.

