## Supplemental Figures for "Selenium protects *Emiliania huxleyi* algae from a bacterial pathogen by modulating oxidative stress"

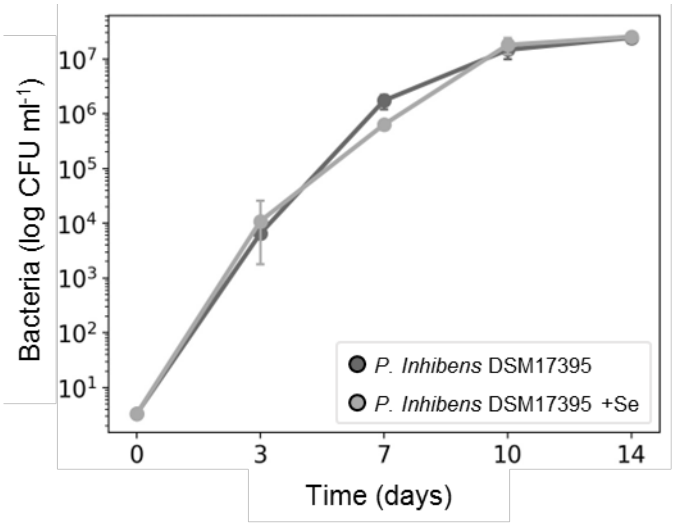


**Figure S1. Growth of *P. inhibens* bacteria in co-cultures with algae.** The growth of the bacterium *P. inhibens* co-cultured with algae along 14 days. Dark gray line – *P. inhibens* grown with algae in co-culture (corresponding to Fig. 1). Light gray line – *P. inhibens* grown with algae in co-culture supplemented with selenium (+Se, corresponding to Fig. 3). Each data point consists of 3 biological replicates, error bars designate ± SD.

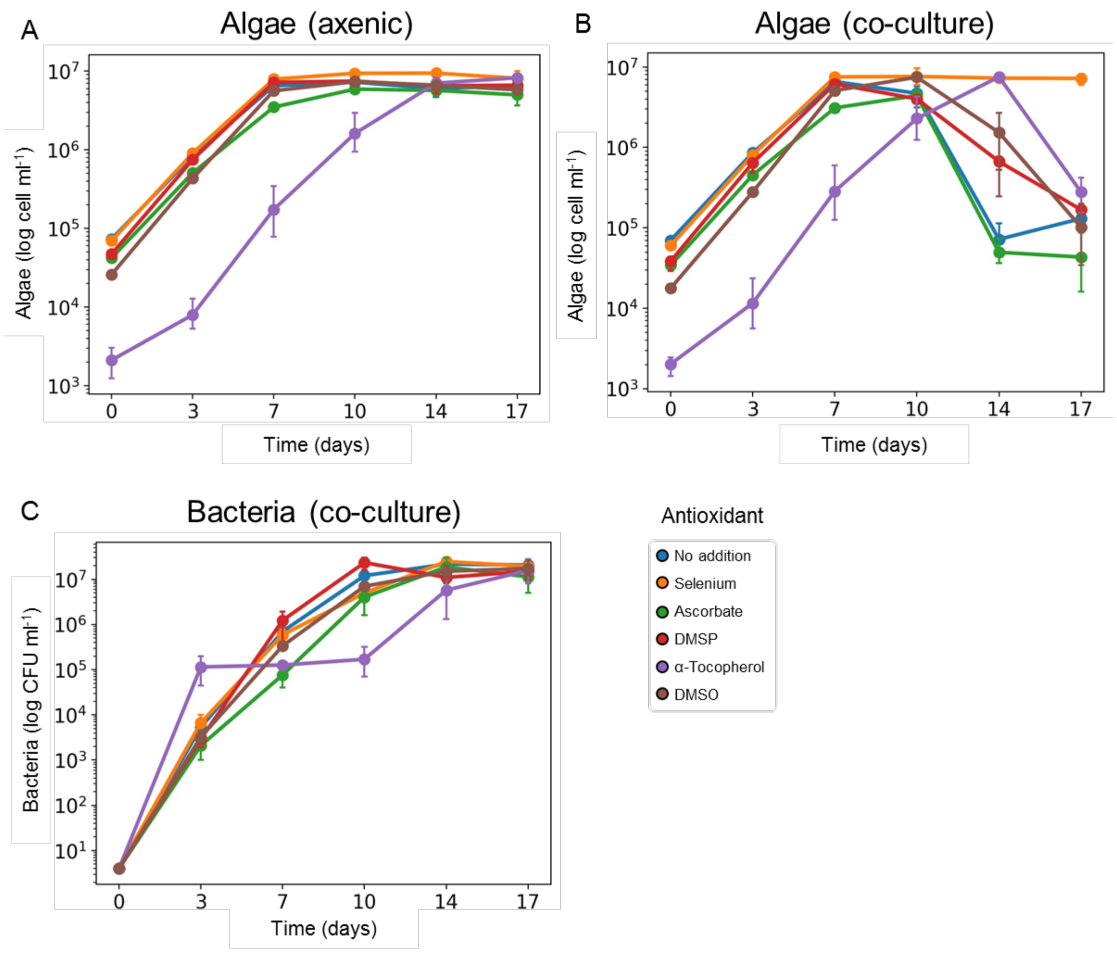


**Figure S2. Growth dynamics of algae and bacteria under different antioxidant treatments.** The algae *E. huxleyi* CCMP2090 was grown either axenically (**A**) or with the bacterium *P. inhibens* DSM17395 (**B**). As indicated, the cultures were supplemented on day 0 of algal growth with different antioxidants – selenium ($H_2SeO_3$, 10 nM), ascorbate (L-Ascorbic acid, 20 μM), DMSP (Dimethylsulfoniopropionate, 20 μM) and α-Tocopherol (20 μM). DMSO (Dimethylsulfoxide, 60 μl) was used as the solvent of α-Tocopherol hence it was supplemented to cultures as a control. Algal cell concentrations were measured along 17 days of growth to account for the influence of the different antioxidants tested. Bacterial growth of *P. inhibens* was monitored along time in co-cultures under the different antioxidant treatments (**C**). Each data point consists of 3 biological replicates, error bars designate ± SD.
